# Genetic connectivity in US Caribbean queen conch (*Aliger gigas*): Implications for management and forensic assignment

**DOI:** 10.64898/2026.09.17.752428

**Authors:** Diana M. Beltrán, Richard Appeldoorn, Carlos Prada

## Abstract

Understanding genetic connectivity among marine populations is central to conservation biology, informing how exploited species are managed, and how marine protected area (MPA) networks are designed. The queen conch (*Aliger gigas*) is a large, heavily exploited Caribbean gastropod listed under CITES Appendix II and provides an ideal system for examining connectivity in species with high dispersal potential. Its extended pelagic larval phase promotes gene flow across broad spatial scales, limiting detectable population structure. Here we used low-coverage whole-genome sequencing (lcWGS) of queen conch sampled from three regions within United States waters, Florida (FL), Puerto Rico (PR), and the US Virgin Islands (VI), to characterize connectivity and to evaluate the implications for management. Using genotype-likelihood approaches, we inferred patterns across principal component analysis, admixture analysis, pairwise F_ST_, contemporary migration, and historical coalescent-based demographic modeling. Genome-wide analyses revealed little structure with individuals from all three regions intermixed in ordination space and similar ancestry profiles, consistent with high gene flow. In contrast, analyses restricted to the upper tail of the F_ST_ distribution recovered geographically concordant structure, resolving FL, PR, and VI as distinguishable groups, with PR individuals showing intermediate, admixed ancestry between FL and VI. Estimates of contemporary migration indicated a partially open system in which 72–98% of each population’s ancestry was locally derived, with substantial bidirectional gene flow between PR and VI, moderate immigration from VI into FL and VI to PR, and negligible direct exchange from FL to either PR or VI. Historical demographic analyses suggest long-term genetic effective population sizes (N_e_) on the order of 2.2 × 10^5^ to 7.6 × 10^5^ individuals and a hierarchical population divergence during the early Pleistocene (∼2 Mya) with ongoing symmetric migration among the three populations, with the strongest historical connectivity between PR and VI with 10 diploid migrants per generation (2Nm = 20.0). Together, these results indicate that queen conch populations across US waters are neither fully panmictic nor completely isolated. Instead, they form a connected population system in which substantial gene flow occurs alongside geographically differentiated genomic variation. These findings argue against treating the US range as either a single management unit or as a set of completely independent stocks. They also highlight the potential importance of the US Virgin Islands within the regional connectivity network and suggest that highly differentiated loci could be useful for developing forensic SNP panels to identify the geographic origin of queen conch products and support enforcement against illegal, unreported, and unregulated fishing.

## INTRODUCTION

Genetic connectivity among marine populations is increasingly recognized as central to conservation biology, shaping how we protect and manage species, design networks of marine protected areas, and gauge the resilience of exploited populations to overfishing and environmental change [1–4]. Among the assumptions long held about the sea is that species with planktonic larvae and high dispersal potential maintain genetically homogeneous populations across broad geographic ranges [2]. Much less straightforward, however, is how often this expectation holds. The accumulation of fine-scale genetic studies has progressively challenged it, revealing population structure at spatial scales far smaller than the theoretical dispersal capacity of larvae would predict [5–7].

Several processes may restrict effective gene flow to a fraction of what dispersal potential alone, often inferred from the movement of particles in physical models, would suggest. Larval philopatry [8,9], oceanographic barriers such as current fronts and gyres [10,11], sweepstakes reproductive success among high-fecundity broadcast spawners [12], and adaptation to local habitats [13] can each, and together, sever the link between how far larvae can travel and how far genes actually move. The result is that marine population structure spans a continuum, from fully open panmixia to virtually closed isolation. Intermediate scenarios, stepping-stone gene flow, geographic clines, and chaotic genetic patchiness, represent the outcomes most often documented for species with complex oceanographic histories [1,2,14,15].

The emergence of genome-wide single-nucleotide polymorphism (SNP) approaches has transformed our ability to resolve this continuum [16,17]. Where microsatellite and allozyme studies often lacked the resolution to detect subtle but ecologically significant differentiation in high-dispersal species, whole-genome datasets now routinely reveal two distinct signatures operating simultaneously: pervasive neutral genomic homogeneity reflecting ongoing larval exchange, and differentiation concentrated in a subset of loci often under divergent natural selection [18–21]. This distinction matters for management. Neutral genetic homogeneity may suggest interchangeable populations that need not be managed as separate units, and fewer and more spaced MPAs [2] Adaptive divergence at selected loci, by contrast, indicates that local populations have evolved distinct physiological or life-history strategies in response to their local environments, and that the exchange of individuals, larvae or gametes across environments carry fitness costs that erode local adaptation and reduce long-term population resilience [22,23]. Understanding the spatial scale of neutral and adaptive variation is therefore essential for designing conservation strategies, delimiting fishing stocks, designing networks of MPAs, and even designing tools to detect poaching and illegal fishing [24].

The queen conch, *Aliger gigas* (Linnaeus 1758; formerly *Lobatus* and *Strombus gigas*) is an ideal model to examine these questions with an urgent conservation priority. This large gastropod is distributed across shallow-water carbonate platforms and seagrass beds of the western Atlantic, from Bermuda and southern Florida to Brazil [25,26], where it occupies a variety of shallow-water habitats including seagrass meadows, coral rubble, algal plains, and sandy substrates, and plays a central herbivory role in these tropical ecosystems [27–29]. Its economic and cultural importance is considerable: queen conch meat has been a dietary staple and export commodity across the Caribbean for centuries, holds deep cultural significance for Caribbean peoples, and underpins a market spanning more than 25 Caribbean nations and territories.

Populations have declined severely across much of the species’ range, driven largely by overfishing and the loss of the critical shallow-water habitats on which the species depends [30,31]. Because conchs are slow-moving and require direct contact to mate, overfishing not only reduces abundance but also depresses reproductive success in depleted areas, driving local densities below the critical threshold for reproductive aggregation and triggering Allee effects that impair recovery even when harvest pressure is relaxed [32–35]. Low population densities of mature organisms have also raised concerns about genetic diversity loss with long-term consequences for adaptive potential [36]. The conservation status of *A. gigas* reflects the severity of these pressures: listed as Commercially Threatened by the IUCN since 1983 [37] and recently as Near Threatened [38], and listed in Appendix II of the Convention on International Trade in Endangered Species (CITES) in 1992, placing its trade under international control [39–41]. In response, Caribbean nations have adopted management measures that have expanded over time and vary considerably among jurisdictions, ranging from minimum size limits and seasonal closures to complete fishing bans in Florida and Bermuda [40].

Effective management of such a transboundary resource hinges on understanding whether queen conch populations function as discrete management units or as a single, demographically connected population. The species’ two-phase life history complicates this question. Adults can be largely sedentary [42,43], although potentially capable of moving up 9+ km given sufficient time and habitat extent [44]. In contrast, long-range dispersal and mixing among populations occurs during the pelagic larval phase, when larvae are transported by oceanic currents over distances that depend on prevailing hydrodynamics. Highly connected populations exhibiting little genetic differentiation effectively constitute a single mixing population that should be managed as one stock, whereas populations displaying strong genetic structure likely represent distinct resources warranting independent management. Despite these observations, the population genetic structure of *A. gigas* remains poorly resolved, with earlier studies producing conflicting conclusions that reflect both the limitations of marker technology and the difficulties of adequate geographic sampling across a species spanning thousands of kilometers of coastline.

Studies employing allozymes and mitochondrial DNA largely concluded that Caribbean queen conch populations constitute a single panmictic or near-panmictic genetic unit [45], an interpretation consistent with the oceanographic connectivity potential of a species whose larvae in culture typically spend approximately three weeks in the water column before settlement [46–48]. A subsequent mtDNA and microsatellite-based study found evidence of an isolation-by-distance pattern among sampled areas, implying that effective gene flow is distance-limited even if gross dispersal potential is high [49,50], while another study detected spatial homogeneity compatible with broad connectivity [51]. These studies were constrained by small numbers of sampling locations, limited numbers of loci, and genetic markers that lacked the resolution to distinguish neutral demographic connectivity from adaptive genetic differentiation, limitations that are now addressable with whole-genome genotyping.

The transition to genome-wide SNP datasets has produced more insights about population structure. Rather than forcing a binary choice between panmixia and isolation, SNP data reveal the coexistence of high connectivity at neutral loci with meaningful differentiation at loci under selection [20]. This pattern is increasingly recognized as a general feature of high-dispersal marine species [21,52,53] with practical consequences for how connectivity is measured and interpreted. Standard multivariate analyses of whole-genome variation, such as PCA and DAPC, weight all loci equally and are therefore dominated by the neutral background, producing results indistinguishable from panmixia even when adaptive divergence is biologically meaningful and spatially consistent [4]. F_ST_-based analyses and AMOVA, being sensitive to variance partitioning rather than pattern recognition, recover the differentiation signal with greater power. Outlier detection methods that identify highly differentiated loci such as F_ST_-based approaches or those that detect loci under divergent selection, including F_ST_ outlier approaches implemented in tools such as BayeScan [54] or OutFLANK [55], isolate the adaptive component of genomic variation and, when used as the basis for multivariate analyses, reveal population structure invisible to genome-wide approaches. This methodological framework has proven effective in species ranging from Atlantic cod [56,57] to bonefish [4,52] and marine invertebrates [21,58], and its application to *A. gigas* offers the potential to resolve persistent uncertainties about population structure in this species.

Here we apply whole-genome SNP analyses to populations of *A. gigas* sampled from three regions within US jurisdictional waters: Florida (FL), Puerto Rico (PR), and the US Virgin Islands (VI). These three locations span the northern and eastern periphery of the queen conch’s range in the Caribbean basin with substantial environmental heterogeneity, from the seasonally variable, turbid, and nutrient-enriched waters of the Florida Keys to the oligotrophic, thermally stable reef systems of the eastern Caribbean. They also represent the primary geographic units under which federal management of US queen conch populations is organized; management has historically been shared between territorial and U.S. federal governments through the Caribbean Fisheries Management Council, with measures including the 1997 closure of the federal Exclusive Economic Zone except around VI (St. Croix), seasonal closures, minimum capture sizes, and bag limits [59,60]. This governance structure makes genetic resolution of connectivity directly relevant to decisions regarding stock identity, harvest allocation, and the design of MPA networks.

We use a combination of genome-wide analyses and F_ST_ approaches to test whether the US populations of *A. gigas* constitute a single panmictic unit or harbor detectable genetic structure, to characterize the connectivity among regions with respect to both neutral and genomic variation tight to high F_ST_ values, and to evaluate the implications of the resulting connectivity patterns for management. We additionally consider the potential of SNPs identified through outlier analyses to serve as the foundation for forensic genotyping tools capable of assigning commercially traded queen conch products to their region of origin, providing a molecular complement to existing frameworks for detecting and deterring illegal, unreported, and unregulated (IUU) fishing, a pervasive threat to *A. gigas* throughout the Caribbean that existing permit-based enforcement mechanisms are insufficient to address [31,40,61,62].

## METHODS

### Sampling, DNA extraction, library preparation and sequencing

We collected tissue samples from queen conch (*Aliger gigas*) from both adults and juveniles across three focal US regions: Florida (FL), Puerto Rico (PR), and the US Virgin Islands (USVI; St. Croix). In Florida, 95 samples were collected from three sites using a non-detrimental survey approach, with a single biopsy punch taken per individual. In Puerto Rico, 50 samples were selected from a larger sample earlier collected between 2014 and 2017 from commercial fishing landings at sites in both the western and eastern parts of the island [60]. In VI (St. Croix), 36 samples were obtained from fisheries independent surveys at seven locations. Within each region, sampling spanned multiple reef systems and localities (Table S1). Tissues were preserved in DNA Shield, RNA later, DMSO, or 95% ethanol at the point of collection.

Genomic DNA was extracted using the E.Z.N.A. Mollusk DNA Kit (Omega Bio-Tek) following the manufacturer’s protocol. DNA concentration was quantified with the AccuBlue Broad Range dsDNA Quantification Kit (Biotium) on a Qubit 2.0 fluorometer and normalized prior to library preparation, and DNA integrity was assessed by 1% agarose gel electrophoresis. Individual dual-indexed libraries were prepared for low-coverage whole-genome sequencing (lcWGS) following a cost-effective protocol [63,64]. Samples coming from FL using the non-invasive method produced very low DNA yields, and many failed to produce viable genomic libraries (26 samples). Fragment size distributions, with a target mean insert size of 350–500 bp, were verified on an Agilent High Sensitivity DNA Bioanalyzer. Libraries were pooled at equimolar concentrations and sequenced with paired-end 150-bp reads on an Illumina NovaSeq platform at a target depth of 5×. To minimize batch effects, samples from different locations were randomly distributed across sequencing runs; final sequencing was carried out across six runs in which our queen conch libraries were multiplexed with libraries from other projects. Following extraction, library preparation, and sequencing QC, the final dataset comprised 69 FL, 50 PR, and 36 USVI individuals (n = 155 total).

### Sequence Filtering and Alignment

Adapters were removed, reads trimmed and quality-filtered with fastp v0.23.4 [65], and per-sample quality metrics were aggregated and visually inspected across all libraries using MultiQC v1.14 [66]. Trimmed reads were mapped using bwa to a de-novo assembled genome using hifiasm [67] with default parameters on publicly available data under project number PRJEB88237. Given that the data come from PacBio libraries this approach allowed us to generate longer contigs and increase mapping accuracy than a regular de-novo assembly using our Illumina short read (PE 150 bp) data. The genome assembly generated a total of 33,439 scaffolds with an average size of 36,022 base pairs and the largest contig of 2,504,132 base pairs. We then indexed the *A. gigas* assembly using bwa-mem v0.7.17 [68] with default settings. Duplicate reads were removed with the Picard v2.25.2 [69].

### Genotype Likelihood Estimation and Dataset Construction

Prior to genotype likelihood estimation, we used ANGSD v0.940 [70] with -doDepth 1 to obtain the genome-wide read-depth distribution and to inform the choice of depth filters. Nineteen individuals with insufficient read depth were excluded from downstream analyses. Genotype likelihoods were then estimated under the Samtools model (-GL 1) [71], retaining reads on the basis of read quality (-uniqueOnly 1, -remove_bads 1, -trim 0, -baq 1) [71], base quality (-minQ 25), linkage (-thin 3000) and mapping quality (-minMapQ 30, -C 50), together with depth filters established from the observed depth distribution (< 5 and > 1500 reads). Polymorphic sites were identified across all individuals using a likelihood-ratio SNP p-value cutoff of 1 × 10⁻⁶. Major and minor alleles were inferred from genotype likelihoods (-doMaf 1, -doMajorMinor 1), sites with a minor allele frequency below 5% were discarded (-minMaf 0.05), and sites with more than two inferred alleles were removed (-rmTriallelic 0.05). Initially we produced a VCF from the ANGSD BCF file and ran OutFLANK [55] and all loci flagged as outliers were excluded to produce putatively neutral SNP sets. We reran ANGSD as stated above removing poorly genotyped samples, and removing all loci flagged as outliers.

In addition, we filtered our dataset to avoid SNPs in physical linkage. Linkage disequilibrium was estimated with ngsLD v1.1.1 [72] using --max_kb_dist 100, LD decay was visualized by plotting LD against physical distance, and correlated variants were removed with the prune_ngsLD.py script. The resulting neutral, LD-pruned dataset was used for both principal component analysis and admixture inference. We generated five different datasets for the different analyses. Dataset I (genome-wide SNPs) comprised the full set of genome-wide SNPs recovered across the largest set of individuals (136) and SNPs (142,418) that passed sample-level filtering. We then applied more stringent SNP and sample filters (--mass-missing > 0.9) to eliminate further missingness in individuals and produced Dataset II. In addition, we pull out the top 5% of SNPs ranked by pairwise F_ST_ (Dataset III, F_ST_ outliers), representing the most strongly differentiated loci. This dataset was also used for principal component analysis and admixture inference to characterize segregated genomic variation. Finally, to corroborate our initial analyses by ANGSD, we produced an independent VCF file by processing our data through the GATK pipeline using the Haplotype Caller algorithm [73] to hard call genotypes across all individuals, producing a multi-sample VCF, excluding SNPs captured by OutFLANK [55] and also thinned with SNPs separated more than 5,000 bases using vcftools [74]. This dataset was also used for an independent principal component analysis and admixture inference (Dataset IV) (Fig. S1).

### Population Structure, Admixture, and pairwise F_ST_

For each genotype-likelihood dataset, we generated an allele-frequency file (-doMaf 1) and a genotype-likelihood file (-GL 1, -doGlf 2) in ANGSD. Individual-level principal component analyses (PCA) were performed from the covariance matrix estimated with -doCov 1 in ANGSD, and eigen-decomposition was carried out in R [70]. To assess the potential influence of missing data on the lcWGS-based PCA, we calculated per-individual missingness and tested its association with individual positions along PC1 (R = 0.02; P > 0.5).

Admixture proportions were estimated from genotype likelihoods with NGSadmix for Datasets II and III, running K = 1 to K = 10 with multiple iterations until convergence [75], defined as a maximum difference of two log-likelihood units among the top replicate runs for each K. Model fit for each value of K was evaluated with evalAdmix v0.962 [75]. Weighted pairwise F_ST_ among sampling regions was estimated in ANGSD from the two-dimensional site frequency spectrum for each pair of regions (realSFS, realSFS F_ST_), using Dataset I as the primary input. These per-site and genome-wide F_ST_ estimates also provided the ranking used to define the top-5% outlier set (Dataset III). As a complementary, hard-call validation, PCA, admixture with cross-validation and F_ST_ analyses were also performed on the GATK-derived Dataset IV using the R package Adegenet [76] and Admixture (Fig. S1), which recovered trends consistent with the genotype-likelihood analyses.

### Contemporary Migration, Effective Population Size and Historical Demography

Contemporary migration rates among FL, PR, and VI were estimated using BayesAss with the SNP-adapted implementation BA3-SNPs [77] applied to a 20k random subset of Dataset II. We ran BA3-SNPs fifteen times with different starting seeds, a burn-in of 2 million and 20 million iterations and combined all runs and reported the average and standard deviation across runs. We assessed convergence of the MCMC and reported directional migration-rate estimates and self-recruitment proportions among the three regions.

We reconstructed the historical demography of FL, PR, and VI using moments [78]. A folded 3-D joint site frequency spectrum was computed de novo with realSFS [70], including all sites and avoiding filters that distort the SFS (Dataset V). Against this spectrum we fit ten three-population models differing in divergence history and migration. Divergence was modeled as either simultaneous (sim_split) or hierarchical (split, in which an ancestral population gives rise to two descendants); for each, we tested no migration, symmetric migration, and asymmetric migration, with gene flow restricted to adjacent populations or occurring among all pairs. Model parameters comprised effective population sizes (nu1, nu2, nu3, and ancestral nuA where applicable), divergence times (T1, T2), and pairwise migration rates (m), ranging in complexity from four parameters (simultaneous divergence, no migration) to thirteen (hierarchical divergence, asymmetric migration among all pairs). We fitted each of the ten demographic models using four successive rounds of optimization in moments, with 10, 20, 30, and 40 replicates per round (100 total per model), progressively increasing the number of optimizer iterations and decreasing the perturbation fold to refine parameter estimates and reduce the risk of convergence on local optima.

The best-supported model was selected by AIC, and its parameters were further refined in three additional fine-tuning rounds (120 replicates) before conversion to absolute units using Nref = θ/(4μL). All optimization replicates used the multinomial likelihood function, which estimates model parameters independently of theta. The replicate yielding the highest log-composite-likelihood was retained as the maximum-likelihood estimate. Parameter estimates were scaled from coalescent units to absolute values assuming a per-generation mutation rate of μ = 2 × 10^−9^ per base, adopted from the oyster *Crassostrea ariakensis* [79,80] in the absence of direct estimates for gastropods, and a generation time of g = 4 years based on reported age at maturity for *Aliger gigas* [81,82].

## RESULTS

To characterize genetic structure among queen conch populations across Florida (FL), Puerto Rico (PR), and the US Virgin Islands (VI), we first examined principal component analyses (PCA) derived from genotype likelihoods (Fig. 1). When all individuals and the complete genome-wide SNP set (Dataset I; 136 individuals, 142,418 SNPs) were analyzed, individuals from the three regions were largely intermingled along the first two principal components, with no clear separation by localities (Fig. 1A). PC1 explained only 1.67% and PC2 1.54% of the total genetic variance, and samples from FL, PR, and VI overlapped extensively. The same pattern persisted in the dataset with fewer individuals (99) and a more stringently filtered SNP panel (Dataset II; 55,671 SNPs), in which individuals again failed to group by locality (Fig. 1B). These analyses indicate that genome-wide variation is shared among regions, consistent with high connectivity and substantial gene flow across the sampled populations.

**Figure 1.**
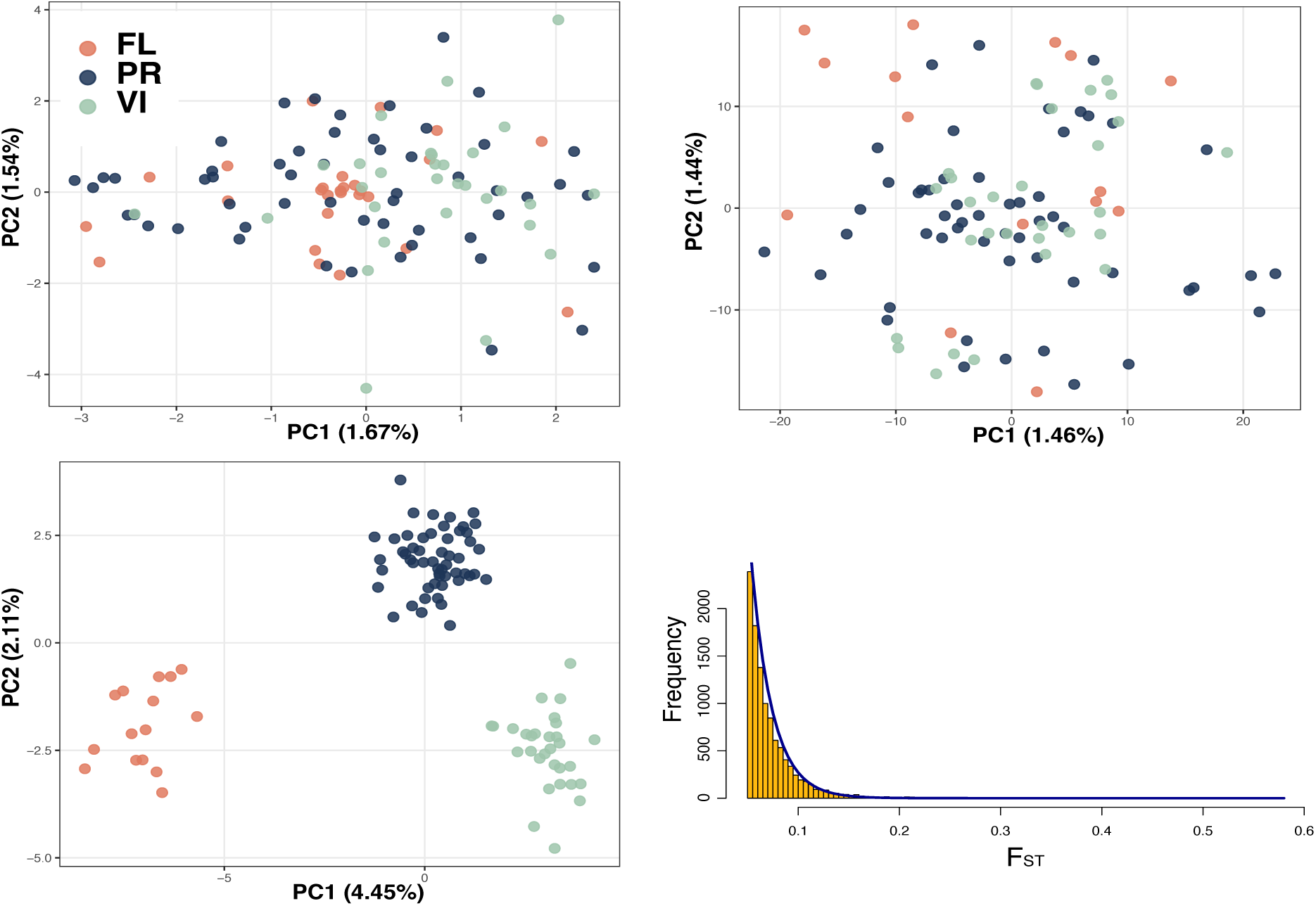
Population structure of queen conch (*Aliger gigas*) from Florida (FL, red), Puerto Rico (PR, blue), and US Virgin Islands (VI, green). (A) PCA of all individuals using the genome-wide SNP set (Dataset I; 136 samples, 142,418 SNPs). (B) PCA of the reduced, lower-missingness neutral SNP set after removal of F_ST_ outliers and LD pruning (Dataset II; 99 samples, 55,671 SNPs). (C) PCA based on the top 5% of SNPs ranked by F_ST_ (Dataset III, 7,120 SNPs). The three regions resolve into clearly distinguishable clusters: FL (red), VI (green), and PR (blue), consistent with PR occupying a central position relative to the two peripheral regions. (D) Genome-wide distribution of per-site F_ST_ values across all polymorphic SNPs, fitted with a smoothed density curve (blue); the distribution is strongly right-skewed, with the great majority of sites showing low differentiation and a long tail of high-F_ST_ loci extending to F_ST_ > 0.5.

A different pattern emerged when the analysis was restricted to the most strongly differentiated loci. Using the top 5% of SNPs ranked by F_ST_ (Dataset III; 7,120 SNPs, see also distribution of F_ST_ values in Fig. 1D), the PCA resolved the three regions into distinguishable groups along the first two axes (Fig. 1C), with FL, PR, and VI occupying separable regions of the ordination space. To further evaluate this pattern, we also used the SNPs identified by OutFLANK as outliers, and recovered a similar pattern of segregation of genetic variation concordant with the origin of the samples (Fig. S2). The contrast between genome-wide gene flow and structure among high F_ST_ loci, indicates that population differentiation is concentrated in a subset of the genome rather than across it. In addition, genome-wide weighted pairwise F_ST_ values among the three regions were low (FL–PR = 0.002, FL–VI = 0.003, PR–VI = 0.0001), consistent with high overall genetic connectivity across the US queen conch range. The pattern is consistent with the weak genome-wide structure that becomes unambiguous when analyses are restricted to high-F_ST_ outlier loci.

We next estimated individual ancestry proportions using NGSadmix on genotype likelihoods (Fig. 2) and, independently, ADMIXTURE on called genotypes (Fig. S1C). Model selection favored K = 2 as the best-supported number of clusters. However, when all SNPs were used, the inferred ancestry clusters showed no correspondence to sampling locality: individuals from FL, PR, and VI carried similar mixed-ancestry profiles, and the cluster assignments did not align with geographic origin (Fig. 2A). This mirrors the PCA results and reinforces the inference of pervasive genome-wide homogeneity among the three regions.

**Figure 2.**
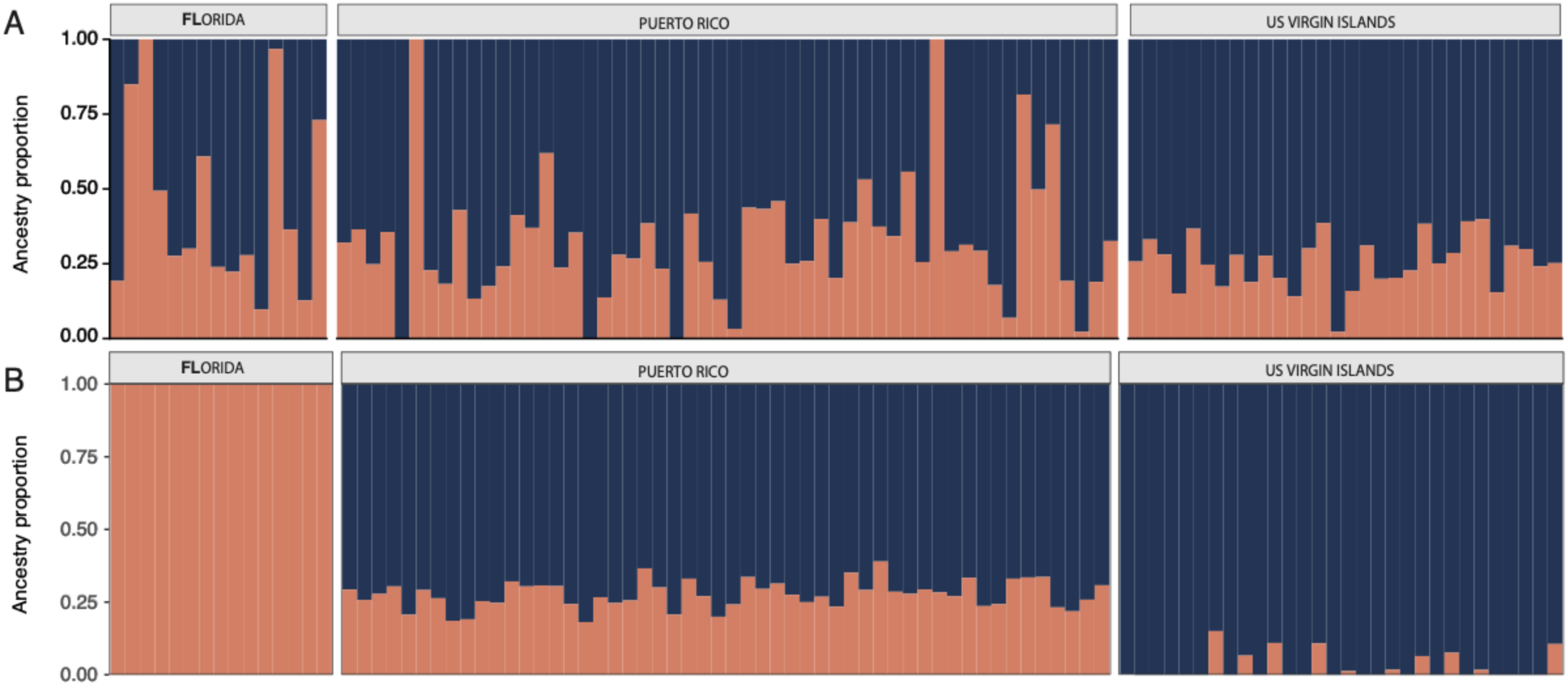
Ancestry proportions for queen conch (*Aliger gigas*) from FL, PR, and VI inferred at K = 2 using NGSadmix, contrasting genome-wide and F_ST_-outlier SNP datasets. Each vertical bar represents one individual, partitioned into ancestry proportions for two inferred clusters (red and blue); individuals are grouped by sampling region. (A) Ancestry proportions estimated from the neutral, LD-pruned genome-wide SNP set (Dataset I). Ancestry assignments show no correspondence with geographic origin. (B) Ancestry proportions estimated from the top 5% of SNPs ranked by F_ST_ (Dataset III). Cluster assignments align with sampling locality with Florida individuals assigned mostly to the red ancestry cluster, VI individuals almost entirely to the blue cluster, and PR individuals with intermediate ancestry. See also Figures S1, and S2 with cross-validation curves for different K values.

When ancestry was instead inferred from the top 5% F_ST_ outlier SNPs or the outlier SNPs, the clusters aligned with sampled localities (Figs. 2B and S2C). Florida individuals were assigned predominantly to one ancestry cluster and US Virgin Islands individuals predominantly to a second, while Puerto Rico individuals showed intermediate, admixed ancestry combining both clusters. This pattern of distinct FL and VI ancestry with admixed PR individuals is consistent with PR occupying a central, intermediate position relative to the two peripheral regions, and parallels the separation recovered in the high F_ST_-based PCA (Fig. 1C). The PCA and admixture analyses suggest that genome-wide markers show no structure and loci in the upper tail of the F_ST_ distribution recover geographical separation, with Puerto Rico intermediate between Florida and the US Virgin Islands.

Contemporary migration rates estimated with BayesAss were highly consistent across independent replicate runs and revealed asymmetric connectivity among the three populations alongside high self-recruitment (Fig. 3). All three populations retained the majority of their members locally each generation, with non-migrant proportions of 0.756 ± 0.009 in FL, 0.716 ± 0.010 in PR, and 0.980 ± 0.000 in the VI. The VI population received negligible immigration from either other region (0.010 from both FL and PR). In contrast, FL and PR both received substantial immigration with markedly asymmetric directionality. FL derived an estimated 0.199 ± 0.013 of its individuals from VI (St. Croix) and 0.046 ± 0.009 from PR, whereas reciprocal flows out of Florida were negligible (< 0.006 into either region). PR received its largest immigrant contribution from VI (St. Croix) (0.279 ± 0.010) while sending very few migrants in return (0.010 into VI). Across all pairs, gene flow was predominantly out of VI (St. Croix) and into both PR and FL, with little or no reciprocal movement, indicating a VI source-biased connectivity into the Northwest of the Caribbean (Fig. 4).

**Figure 3.**
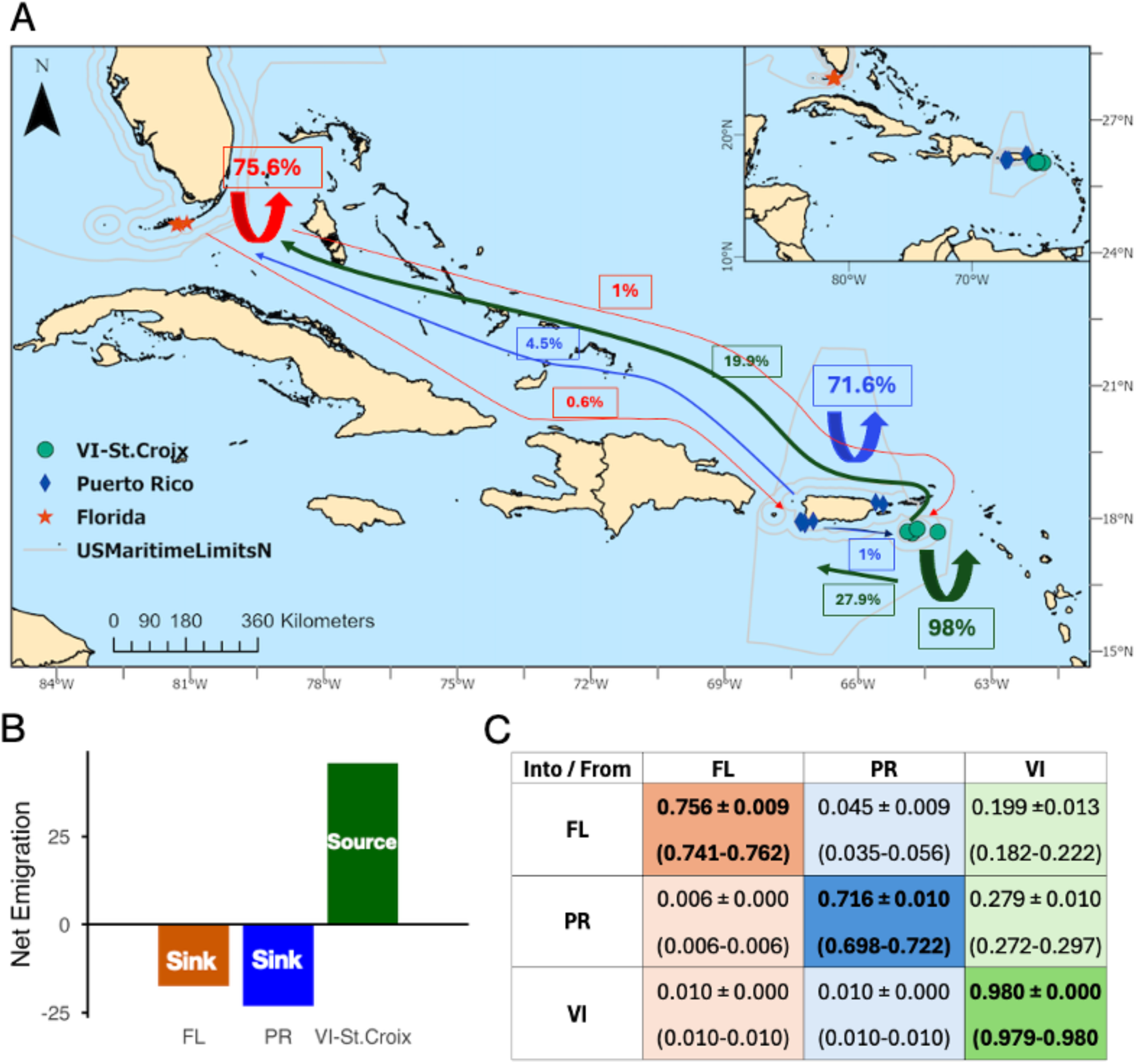
**A**. Contemporary migration among FL (red stars), PR (blue diamonds), and VI (green circles). Thin blue lines delineate US Exclusive Economic Zone boundaries. Curved arrows centered on each region indicate the proportion of individuals that are local in origin (self-recruitment): 75.6% in FL (red), 71.6% in PR (blue), and 98.0% in VI (St. Croix) (green). Straight arrows between regions indicate the directional per-generation migration rates, with the color of each arrow matching the source population and arrow thickness scaled to migration rate. **B.** Asymmetric migration inferred among FL, PR and VI. Values represent net migration rates (total emigration minus total immigration) estimated for each population. Positive values indicate populations acting as net sources of migrants, whereas negative values indicate net sinks. **C.** Contemporary migration rates among FL, PR, and VI combined across 15 replicate runs with different seeds (mean ± SD; min–max range). Rows are the receiving population; columns are the source. Diagonal bold values are self-recruitment; off-diagonal values are the per-generation fraction of the population derived from each source.

Demographic inference using moments supported a model of hierarchical population divergence with ongoing symmetric migration among all three populations (Table S2). The best model, split_sym_mig_all, describes an initial divergence separating one lineage (FL) from the ancestor of the remaining two populations (PR and VI), followed shortly thereafter by a second split. The inferred ancestral effective population size was large (Nref = 2.2 × 10^5^), and descendant populations remained comparatively large, with PR showing the largest effective population size (N_e_ = 7.6 × 10^5^), followed by VI (N_e_ = 3.5 × 10^5^) and FL (N_e_ = 2.2 × 10^5^). The two divergence events were closely spaced in time, occurring approximately 2.0 and 1.9 million years ago, respectively, suggesting that diversification began during the early Pleistocene. Migration estimates suggest gene flow between the first-diverging lineage and the ancestor of the other two populations was negligible during the earliest divergence phase (2Nm = 0.09), whereas migration among the contemporary FL, PR, and VI populations was high and symmetric. Connectivity was strongest between PR and VI (2Nm = 20.0), followed by FL–VI (2Nm = 18.7) and FL–PR (2Nm = 16.7), indicating substantial historical and ongoing demographic connectivity across the region. The strong support for a model incorporating migration among all population pairs, together with the poor fit of more restrictive isolation models (Table S2), suggests that gene flow has played a key role in shaping patterns of genetic variation. The strong connectivity between PR and VI is also consistent with the BayesAss results, which identified contemporary exchange between these eastern Caribbean populations, supporting a scenario in which both historical and contemporary gene flow contribute to the observed population structure.

## DISCUSSION

The population genetic structure of *Aliger gigas* across Florida, Puerto Rico, and the US Virgin Islands shows a pattern of a marine species with high dispersal with weak but detectable differentiation that is masked at the genome-wide level by gene flow, yet becomes unambiguous when analyses are focused on loci with high F_ST_ values. The pattern of panmixia-like in neutral genomic variation alongside structure in differentiated loci, has implications for how we interpret connectivity in this commercially and ecologically important gastropod, for how MPA networks should be designed to protect it, and the best way to design SNP-based tools to capture illegal fishing.

### Weak but significant genome-wide structure

The results from pairwise F_ST_ analyses suggest population differentiation among FL, PR, and VI, consistent with a partially open metapopulation system. However, the magnitude of this differentiation is modest. In species with large effective population sizes and extended pelagic larval durations, such as the queen conch, even small F_ST_ values can be statistically significant while reflecting substantial gene flow. *A. gigas* larvae spend approximately three weeks in the water column before settlement [61], and oceanographic modeling of Caribbean circulation suggests that larvae can potentially disperse hundreds of kilometers during this period [83,84] . The relatively high residency values recovered by BayesAss (72%–98%) for all three populations, are consistent with a partially closed population in which the majority of recruitment is self-seeded, but a substantial number of individuals in each generation are migrants from elsewhere, particularly from neighboring populations.

The failure of whole-genome PCA to recover population structure is not surprising and does not indicate an absence of differentiation. When gene flow is high relative to genetic drift, the bulk of genomic variation is shared among populations, and multivariate ordination methods that weight all SNPs equally will be dominated by neutral variation that carries no population signal. This is a well-documented limitation of PCA-based approaches in high-gene-flow marine systems [18,19] and should not be interpreted as evidence that populations are interchangeable. Rather, the signature of population identity in highly connected populations is not in overall allele frequency differences across the genome, but in a small subset of loci that likely respond to local environments.

### Population structure across high F_ST_ loci

A key finding of this study is the emergence of population structure when analyses are restricted to SNPs with high F_ST_ values. These loci produce well-separated population clusters corresponding to FL, PR, and VI, a pattern entirely absent from the neutral genomic background. This divergence between neutral and likely adaptive genomic patterns is often referred as divergence with gene flow and is increasingly recognized as a common feature of marine species whose populations are connected by larval dispersal but differentiated by local adaptation [20,21,85].

The implication is that while neutral alleles are homogenized across the range by larval exchange, a subset of loci likely under divergent natural selection maintain population-specific allele frequencies despite that gene flow. In *A. gigas*, the environmental differences among FL, PR, and VI are substantial. Florida populations inhabit seagrass beds and hardbottom habitats along the Florida Keys, where temperature seasonality is more pronounced, nutrient regimes differ from oligotrophic Caribbean waters [86,87] and hydrodynamic exposure varies markedly from the wave-swept exposed reefs typical of VI habitats [43]. Puerto Rico occupies an intermediate oceanographic position, exposed to both Atlantic and Caribbean water masses depending on season and location [88]. These environmental contrasts provide ample opportunity for divergent selection to act on loci related to thermal tolerance, feeding strategy, or shell morphology, processes that could maintain population-specific allele frequencies at a subset of loci even against the homogenizing force of ongoing gene flow.

The identity of the high F_ST_ loci and the biological processes they underlie remain to be fully characterized, but the pattern itself is consistent with ecological divergence operating at spatial scales within the dispersal potential of the species. This is precisely the scenario described by models of local adaptation with gene flow [89], in which adaptation is encoded in a number of loci of small effect or many of small polygenic nature that are more resistant to the allele frequency erosion imposed by migration [90].

### Connectivity asymmetry and VI (St. Croix) as a net source

Contemporary migration estimates depict a partially open system in which each population is largely self-sustaining but linked by asymmetric, directional gene flow. Self-recruitment of 72–98% across the three regions is consistent with a pelagic larval phase with substantial but limited exchange (Fig. 3A). The pronounced asymmetry with immigration into FL and PR but negligible export from either, against an almost entirely closed VI, suggests that realized larval transport is directional rather than symmetric, likely reflecting prevailing Caribbean current pathways that move larvae preferentially from the eastern portions of the range. The strong VI-to-PR and VI-to-FL contributions, combined with the near-absence of return flow, are consistent with VI and eastern Caribbean functioning as upstream sources whose reproductive output subsidizes downstream northwestern populations (Fig. 3C). This implies that Florida and Puerto Rico may be demographically dependent in part on upstream recruitment, so that depletion of eastern Caribbean sources could erode recovery potential in the receiving populations even where local habitat remains intact. This source–sink signal underscores that management of US queen conch should not treat these regions as fully independent stocks, since the persistence of the receiving populations appears linked to the protection of their upstream sources from the Lesser Antilles. This signal should become evident in a Caribbean-wide analysis.

The pattern documented here of limited neutral differentiation alongside structure at high F_ST_ loci is not unique to US queen conch. A parallel case in bonefish (*Albula vulpes*) and queen conch in the Bahamian Basin [4] similarly recovered near-panmictic patterns from standard analyses but clear, geographically coherent structure with a subset of outlier loci, mirroring results in other marine species [52,53]. This convergence across distinct species and locations suggests that the disconnect between neutral panmixia and adaptive divergence is a general feature of high-dispersal Caribbean marine organisms [91,92].

Our results also reveal source–sink dynamics, with the US Virgin Islands functioning as a connectivity hub supplying migrants to both Florida and Puerto Rico, broadly consistent with Caribbean circulation patterns [93]. This contrasts with Bahamian queen conch [4], which appear demographically independent with strong signals of isolation despite broadly similar larval durations, underscoring that oceanographic connectivity potential does not translate uniformly into realized gene flow and that species-specific larval behavior, settlement ecology, and habitat specialization can produce divergent demographic outcomes. The pattern documented here for *A. gigas* across FL, PR, and VI may reflect the greater inter-site distances, less constrained oceanographic pathways, and more pronounced environmental contrasts.

### Historical effective population sizes

The large long-term effective population sizes recovered by demographic modeling (Ne = 2.2–7.6 × 10^5^) are consistent with the expectation that queen conch, like other broadcast-spawning marine invertebrates, historically maintained large populations across the Caribbean. However, effective population sizes are often orders of magnitude smaller than census size due to high variance in reproductive success [12]. The Ne values reported here should therefore not be interpreted as current census counts but rather as long-term averages over the coalescent timescale, reflecting the genetic footprint of populations that were historically far larger than they are today. It also suggests that the contemporary population decline, driven by overharvesting throughout the Caribbean [94] may have not yet significantly affected the species’ historical effective population size.

### Early Pleistocene divergence with secondary connectivity

The divergence (∼2.0 and 1.9 Mya) occurred during the early Pleistocene, a period of glacial–interglacial sea-level oscillation that repeatedly restructured Caribbean shallow-water habitats and has been implicated in the diversification of other marine taxa in the region [95,96]. The near-zero ancestral migration (2Nm = 0.09) during the earliest divergence suggests that the split between FL and the PR–VI ancestor occurred under isolation, potentially driven by habitat fragmentation during a glacial lowstand. The initial isolation did not persist given the high contemporary migration estimates (2Nm = 16.7–20.0, corresponding to approximately 8.4–10 effective diploid migrants per generation among all population pairs), which far exceed the theoretical threshold of one migrant per generation required to prevent differentiation by drift alone [97]. The ancient divergence followed by gene flow is consistent with a metapopulation maintained by larval connectivity across the wider Caribbean, in which historical separation has not produced a strong present-day genetic structure, concordant with the low pairwise F_ST_ values observed across the three populations.

### Implications for MPA network design

The combination of high connectivity and differentiation at high-F_ST_ loci has implications for MPA design. Traditional approaches in high-dispersal species focus on spacing networks within the larval dispersal kernel so that protected sources can seed adjacent fished areas [2,98] . For queen conch across the US range, the data support this framework with substantial gene flow, between PR and VI, with no evidence of the deep phylogeographic isolation that would require treating these populations as independent units.

However, the existence of differentiation at a subset of loci complicates this picture. If high-F_ST_ loci encode local adaptations to distinct thermal and hydrodynamic environments of Florida versus Caribbean reef systems, indiscriminate translocation across these boundaries for restoration could introduce maladapted genotypes, reduce local fitness, and disrupt co-adapted gene complexes [99,100]. This concern parallels outbreeding depression documented in other marine invertebrates transplanted across oceanographic boundaries [22] and depth-associated adaptive divergence in Caribbean octocorals that maintain reproductive isolation against high gene flow [21,101]. The conservation priority suggested by these data is maintaining the connectivity among PR and VI that sustains gene flow throughout the system; if VI functions as a regional source, its protection should be treated as a network priority.

### The queen conch as a model for dispersal–connectivity inconsistencies

The disconnect between *A. gigas* larval dispersal potential and the modest but detectable differentiation documented here is consistent with the “sweepstakes reproductive success” or “many migrants but few successful settlers” phenomenon [12]. Even species with long-distance dispersal show population structure when larval exchange falls short of dispersal potential, because most far-traveling larvae fail to survive settlement or compete successfully for post-settlement habitat [102]. The combination of high dispersal with site-specific selection implied by our data would produce the pattern observed with modest F_ST_ at neutral loci, asymmetric migration rates, and differentiation at high-F_ST_ loci.

This interpretation suggests that managing queen conch as a panmictic Caribbean population, as some fisheries frameworks have proposed, would underestimate the ecological significance of local differentiation and the population-specific components of recovery. Conversely, treating FL, PR, and VI as completely isolated stocks would overstate genetic independence and miss the conservation benefit of cross-regional connectivity for demographic resilience.

### Genomic tools to capture illegal, unreported, and unregulated fishing

The high-F_ST_ SNPs identified here provide a mechanism for developing enforcement tools against illegal, unreported, and unregulated (IUU) fishing, which threatens food security and has contributed to the species’ decline and its listing under CITES Appendix II since 1992 [61]. IUU fishing also complicates fisheries management [40]; Jamaica, for example, subtracts IUU estimates from its catch quotas before allocating shares to the fishery [103].

Assigning confiscated products to a region of origin would complement existing regulatory frameworks. As shown in Atlantic cod, gene-associated SNPs can assign marine fish to their population with high precision [56], and because these loci are diverging rather than drifting, they should maintain temporally stable, population-specific allele frequencies, a critical requirement for forensic tools that must remain reliable without recalibration.

F_ST_-ranked SNP panels like the ones we described here could let enforcement agencies determine whether a confiscated catch originated from US or other Caribbean waters. Developing one would require sampling across the wider Caribbean and building a database of populations common in international trade. Combined with the CITES framework and the US Seafood Import Monitoring Program, such a tool would provide independent molecular verification of mislabeled or fraudulent shipments that current systems cannot reliably identify.

## ACKNOWLEDGMENTS

We thank Martha Prada, the Caribbean Fisheries Management Council and the Gulf and Caribbean Fisheries Institute for their invaluable logistical support. Samples from Florida were collected by Gabriel Delgado from the Fish and Wildlife Research Institute at the Florida Fish and Wildlife Conservation Commission. We also thank Sennai Habtes, Eva M. Collazo Montanez and Jesus Rivera Hernandez of the USVI Department of Planning and Natural Resources for facilitating sample collection in St. Croix, USVI. Andrés Maldonado and Edgardo Ojeda facilitated sample collection in Puerto Rico. We extend special thanks to Brittany Hanley, Elizabeth Welch, Luis Perez, and Willow Dunster at URI for their assistance with laboratory work and library preparation. We performed sequencing at the Oklahoma Medical Research Foundation facility. This work was supported by Umoja Parent Grant S1-32QTL-000033, US-NOAA 2020, CITES Project No. S-650 and GCFI environmental Network, CITES project No. S-686.

## DATA AVAILABILITY

All raw sequence data in this study have been deposited in the NCBI Sequence Read Archive (SRA) under BioProject accession [TBD]. The *Aliger gigas* reference genome assembled and used for read mapping was generated from publicly available PacBio data under ENA project accession PRJEB88237. Sample metadata, the folded joint site-frequency spectrum, and the scripts used for filtering, population-structure analysis, migration estimation, and demographic modeling are available at [Zenodo DOI].

## AUTHOR CONTRIBUTIONS

D.M.B. and C.P. conceived and designed the study. D.M.B. and R.A. collected samples and coordinated field logistics. D.M.B. performed DNA extractions, prepared sequencing libraries, and carried out bioinformatic and population-genomic analyses. C.P. supervised the analyses and contributed with analysis. R.A. provided fisheries and management context. D.M.B. and C.P. wrote the manuscript with input from R.A. All authors read and approved the final manuscript.

## Notes

https://www.ncbi.nlm.nih.gov/sra?linkname=bioproject_sra_all&from_uid=1248367

### Competing Interest Statement

The authors have declared no competing interest.

