## Supplementary figures and tables for "Genetic connectivity in US Caribbean queen conch (*Aliger gigas*): Implications for management and forensic assignment"

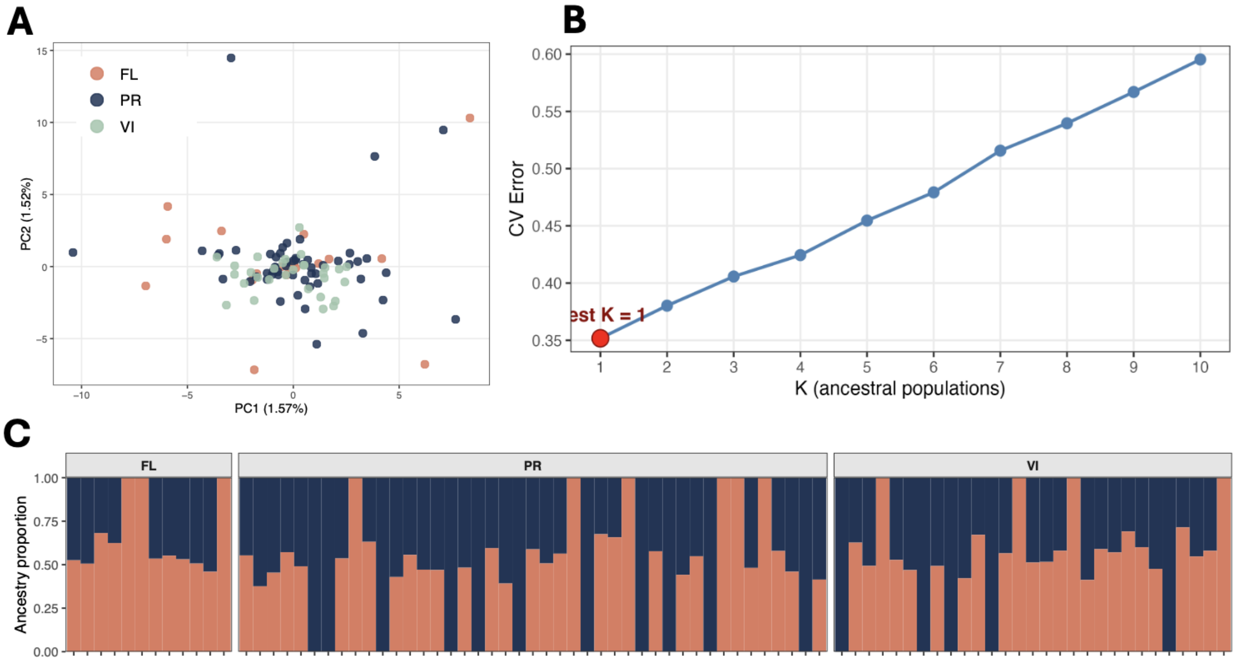


Figure S1. Population structure of queen conch (*Aliger gigas*) across the three US regions, including Florida (FL, red), Puerto Rico (PR, blue), and the US Virgin Islands (VI, green) and inferred from SNPs derived from the GATK pipeline (Dataset IV). (A) Principal component analysis of individual genotypes. (B) Cross-validation (CV) error for ADMIXTURE runs across K = 1 to 10 ancestral populations; CV error is minimized at K = 2 (red point), identifying two ancestral clusters as the best-supported model. (C) Individual ancestry proportions at K = 2, with each vertical bar representing one individual partitioned into the two inferred ancestral clusters (red and blue) and grouped by sampling region.

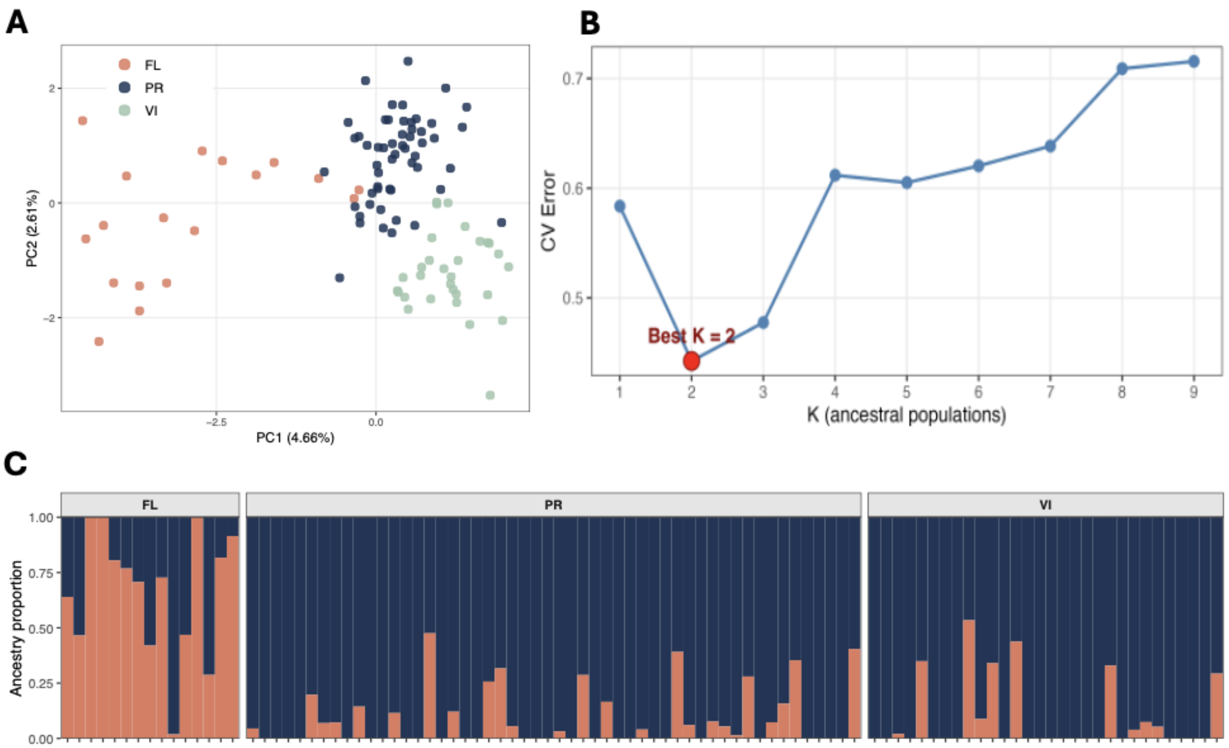


Figure S2. Population structure of queen conch (*Aliger gigas*) across the three US regions, including Florida (FL, red), Puerto Rico (PR, blue), and the US Virgin Islands (VI, green) and inferred from SNPs under selection according to OutFLANK. (A) Principal component analysis of individual genotypes. (B) Cross-validation (CV) error for ADMIXTURE runs across K = 1 to 9 ancestral populations; CV error is minimized at K = 2 (red point), identifying two ancestral clusters as the best-supported model. (C) Individual ancestry proportions at K = 2, with each vertical bar representing one individual partitioned into the two inferred ancestral clusters (red and blue) and grouped by sampling region.

Table S1. Collection localities for queen conch (*Aliger gigas*) sampled across Puerto Rico, US Virgin Islands, and Florida. For each collection, the table reports geographic coordinates, region, collection date, named collecting area (where available), and the number of individuals sampled. Puerto Rico samples were collected in 2014 and 2017; St. Croix and Florida samples were collected in 2023 and 2025. A total of 181 individuals were collected across 21 collections.

| | LONGITUDE | LATITUDE | LOCATION | COLLECTION DATE | AREA | INDIVIDUALS |
|---|---|---|---|---|---|---|
| 1 | -67.00781 | 17.94300 | Puerto Rico | 23-Jan-14 | La Parguera | 2 |
| 2 | -67.18532 | 17.88835 | Puerto Rico | 21-Feb-14 | Cabo Rojo-I | 3 |
| 3 | -67.31075 | 17.93091 | Puerto Rico | 4-Mar-14 | Cabo Rojo-II | 2 |
| 4 | -67.30705 | 17.93898 | Puerto Rico | 6-Mar-14 | Cabo Rojo-III | 1 |
| 5 | -67.28980 | 17.90896 | Puerto Rico | 26-Mar-14 | Cabo Rojo-IV | 1 |
| 6 | -67.19851 | 17.88658 | Puerto Rico | 24-Apr-14 | Cabo Rojo-V | 4 |
| 7 | -67.26101 | 17.93351 | Puerto Rico | 13-May-14 | Cabo Rojo-VI | 2 |
| 8 | -65.60417 | 18.36350 | Puerto Rico | 12-Jun-14 | Fajardo-I | 15 |
| 9 | -65.43750 | 18.32083 | Puerto Rico | 27-Jun-14 | Fajardo-II | 17 |
| 10 | -67.30333 | 17.92963 | Puerto Rico | 27-Jun-17 | Cabo Rojo-VIII | 2 |
| 11 | -67.30333 | 17.92963 | Puerto Rico | 27-Jun-17 | Cabo Rojo-IX | 1 |
| 12 | -64.2076 | 17.69554 | St. Croix | 10-Jan-25 | Point 1 | 5 |
| 13 | -64.77822 | 17.67578 | St. Croix | 29-Jan-25 | Point 2 | 5 |
| 14 | -64.78447 | 17.66784 | St. Croix | 30-Jan-25 | Point 3 | 5 |
| 15 | -64.886385 | 17.704112 | St. Croix | 19-Feb-25 | Point 4 | 2 |
| 16 | -64.885318 | 17.717379 | St. Croix | 19-Feb-25 | Point 5 | 3 |
| 17 | -64.885318 | 17.717379 | St. Croix | 21-Feb-25 | Point 6 | 6 |
| 18 | Unknow | Unknow | St. Croix | Unknow | Unknow | 10 |
| 19 | -81.3126 | 24.64097 | Florida | 23-Oct-23 | Scout Key | 35 |
| 20 | -81.07343 | 24.68713 | Florida | 29-Jan-25 | East Sisters | 17 |
| 21 | -81.24434 | 24.6655 | Florida | 2-Jul-25 | Pirates | 43 |
| | | | | | **TOTAL** | 181 |

Table S2. Comparison of ten three-population demographic models fitted to the folded joint site-frequency spectrum of queen conch (*Aliger gigas*) from Florida (FL), Puerto Rico (PR), and U.S. Virgin Islands (VI) using moments. Models are ranked by Akaike Information Criterion (AIC; lower values indicate better fit). ΔAIC is the difference in AIC relative to the best-supported model (split_sym_mig_all). Log-likelihood, chi-squared goodness-of-fit, and the optimal value of theta (θ) are reported for the best replicate of each model across four rounds of optimization (100 replicates per model). Models prefixed "sim_split" assume simultaneous three-way divergence; models prefixed "split" assume sequential divergence in which one population diverges first, followed by a second split.

| Model | Replicate | log-likelihood | AIC | chi-squared | theta | DELTA-AIC |
|---|---|---|---|---|---|---|
| **split_sym_mig_all** | Round_2_Replicate_14 | -91887.23 | 183794.46 | 1.51E+25 | 27906.79 | - |
| **split_nomig** | Round_4_Replicate_15 | -98383.36 | 196780.72 | 6.17E+22 | 29007.56 | 12986.26 |
| **sim_split_asym_mig_all** | Round_2_Replicate_4 | -108962.73 | 217945.46 | 3.10E+31 | 11408.51 | 34151 |
| **sim_split_no_mig** | Round_4_Replicate_33 | -114171.48 | 228350.96 | 4.48E+27 | 60159.02 | 44556.5 |
| **split_symmig_adjacent** | Round_4_Replicate_10 | -116254.66 | 232525.32 | 1.07E+20 | 2441.36 | 48730.86 |
| **sim_split_sym_mig_all** | Round_4_Replicate_1 | -118536.79 | 237087.58 | 6.15E+21 | 29810.66 | 53293.12 |
| **split_asymmig_adjacent** | Round_4_Replicate_1 | -119124.31 | 238266.62 | 1.72E+25 | 9333.61 | 54472.16 |
| **split_asym_mig_all** | Round_3_Replicate_1 | -168998.48 | 338022.96 | 3.6023E+11 | 15390.54 | 154228.5 |
| **sim_split_sym_mig_adjacent** | Round_2_Replicate_13 | -173886.98 | 347785.96 | 1.16E+22 | 7073.07 | 163991.5 |
| **sim_split_asym_mig_adjacent** | Round_3_Replicate_4 | -267785.57 | 535587.14 | 21855860.8 | 8833.43 | 351792.68 |

Table S3. Maximum-likelihood parameter estimates from the best-supported demographic model (split_sym_mig_all) for queen conch (*Aliger gigas*) across Florida (FL), Puerto Rico (PR), and the US Virgin Islands (VI), inferred with *moments*. The left columns report parameters in coalescent (scaled) units as estimated by the model; the right columns report their conversion to conventional ecological units. Effective population sizes (Ne) are shown for each contemporary population and for the ancestral populations, and were obtained by scaling the relative sizes (nu1, nu2, nu3, nuA) to the reference effective size Nref, derived from θ and the effective sequence length (L = 12,013,208 bp) assuming a per-generation mutation rate of $\mu = 2 \times 10^{-9}$ per base. Divergence times (T1, T2), scaled by 2Nref generations, were converted to years assuming a generation time of 4 years, placing the first split at ~2.05 Mya and the second at ~1.95 Mya. Migration parameters are reported both as the model-scaled rates (m) and as the number of migrants exchanged per generation (2Nm): mA denotes ancestral migration between the first-diverging lineage (FL) and the ancestor of PR+VI (2Nm = 0.09), while m12 (FL↔PR), m23 (PR↔VI), and m13 (FL↔VI) denote symmetric contemporary migration among the three populations (2Nm = 16.7, 20.0, and 18.7, respectively).

| **Model Parameter** | **Value** | **Scaled Parameter** | **Value** |
|---|---|---|---|
| nu1 (FL) | 0.995034 | Nref | 221,000 |
| nuA (ancestral PR+VI) | 3.050622 | N_FL | 220,000 |
| nu2 (PR) | 3.451885 | N_ancestral | 674,000 |
| nu3 (VI) | 1.568261 | N_PR | 763,000 |
| mA | 0.086124 | N_VI | 347,000 |
| m12 (FL↔PR) | 16.666105 | First split | 2.05 Mya |
| m23 (PR↔VI) | 19.977874 | Second split | 1.95 Mya |
| m13 (FL↔VI) | 18.683783 | mA (2Nm) | 0.086 |
| T1 | 0.058936 (first epoch) | m12 (2Nm) | 16.67 |
| T2 | 1.100824 (second epoch) | m23 (2Nm) | 19.98 |
| | | m13 (2Nm) | 18.68 |
| | | L values12,013,208 | |
